# Gut microbiota and metabolites drive chronic sickle cell disease pain

**DOI:** 10.1101/2023.04.25.538342

**Authors:** Katelyn E. Sadler, Samantha N. Atkinson, Vanessa L. Ehlers, Tyler B. Waltz, Michael Hayward, Dianise M. Rodríguez García, Nita H. Salzman, Cheryl L. Stucky, Amanda M. Brandow

## Abstract

Pain is a debilitating symptom and leading reason for hospitalization of individuals with sickle cell disease. Chronic sickle cell pain is poorly managed because the biological basis is not fully understood. Using transgenic sickle cell mice and fecal material transplant, we determined that the gut microbiome drives persistent sickle cell pain. In parallel patient and mouse analyses, we identified bilirubin as one metabolite that induces sickle cell pain by altering vagus nerve activity. Furthermore, we determined that decreased abundance of the gut bacteria *Akkermansia mucinophila* is a critical driver of chronic sickle cell pain. These experiments demonstrate that the sickle cell gut microbiome drives chronic widespread pain and identify bacterial species and metabolites that should be targeted for chronic sickle cell disease pain management.

**One-Sentence Summary:** Gut microbes and metabolites drive chronic sickle cell disease pain by altering vagus nerve activity.

## Main text

Individuals with sickle cell disease (SCD), a β-globin hemoglobinopathy and the most common genetic blood disorder in the world, live with widespread chronic pain. Because the biological basis of sickle cell pain is poorly understood, few effective non-opioid based treatment options exist. Disease modifying therapies such as hydroxyurea and curative therapy such as hematopoietic stem cell transplant decrease pain reports *(1)*, thus suggesting that ongoing red blood cell sickling, vaso-occlusion, tissue hypoxia, and hemolysis contribute, in part, to SCD pain. These physiological processes and their inflammatory metabolic products can disrupt the environments in which gut bacteria reside. Further, daily administration of prophylactic penicillin from birth through six years – a standard-of-care practice to prevent life-threatening pneumococcal sepsis in young children with SCD – could further alter the intestinal microbiota. In support of this concept, recent 16S rRNA gene sequencing studies revealed differences in the number and types of bacteria found in fecal material from individuals with SCD relative to healthy, age-, race-, and gender-matched controls *(2,3)*. However, it is unclear whether this gut dysbiosis contributes to chronic SCD pain or is simply a result of underlying disease pathology. Here, we used transgenic SCD mice and plasma samples from individuals with SCD to determine the extent to and mechanisms through which the gut microbiome contributes to chronic SCD pain.

### SCD gut contents drive persistent pain by altering vagal afferent excitability

To first determine if manipulation of the gut microbiome alters chronic SCD pain, mechanical pain thresholds of mice expressing normal (Townes AA; hemoglobin control) or sickle (Townes SS; SCD) human β-globin *(4)* were tested after animals drank water supplemented with the probiotic VSL#3 or penicillin, an antibiotic that is taken daily by young children *(5)* and many adults with SCD *(6,7)*. Antibiotic administration increased the mechanical sensitivity of both SCD and control animals, whereas probiotic administration completely reversed the mechanical allodynia observed in SCD mice (Fig. 1A).

**Fig. 1.**
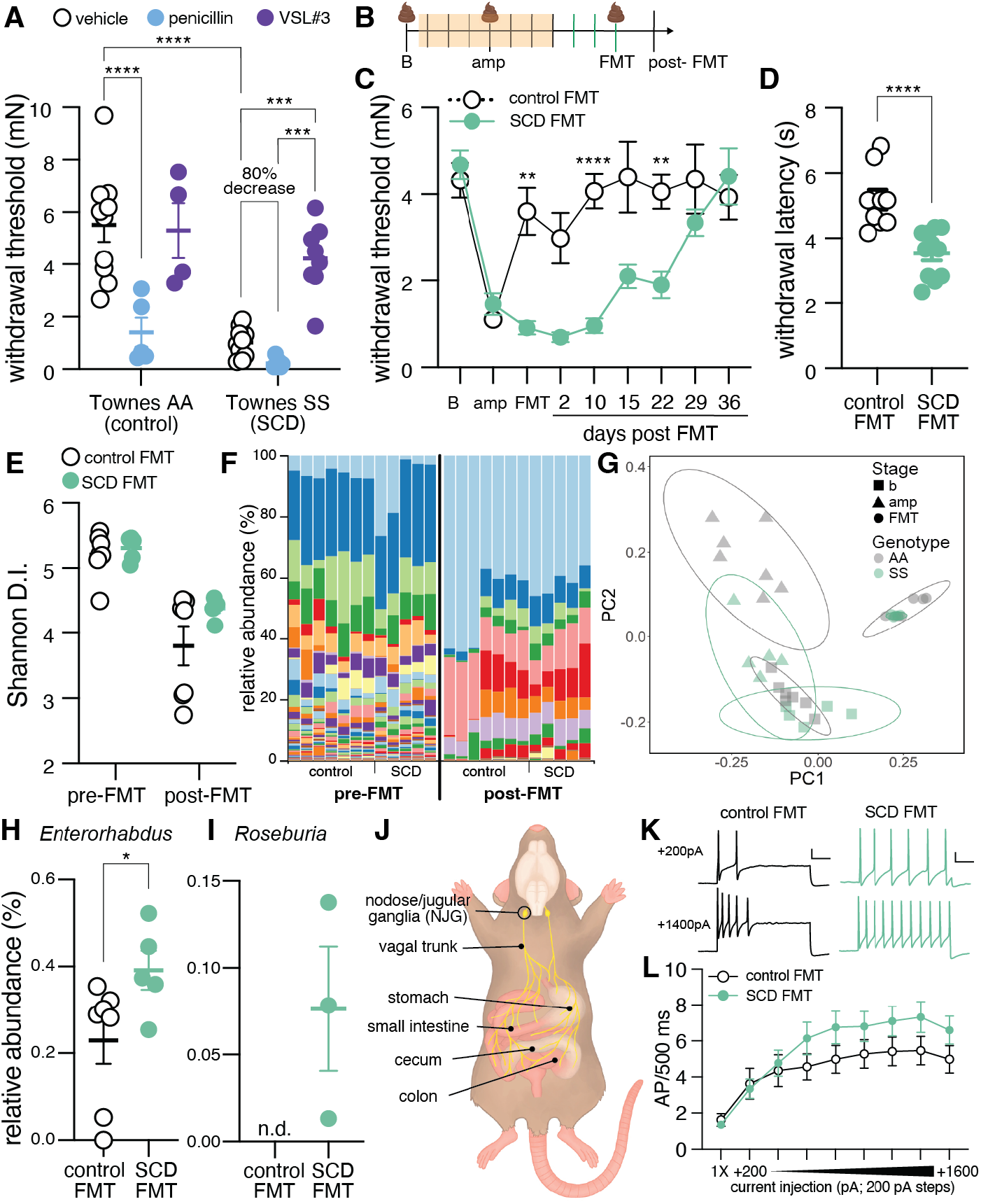
SCD gut contents drive persistent pain. **(A)** Hindpaw mechanical withdrawal thresholds of Townes AA (control) and Townes SS (SCD) mice following 7 days of *ad libidum* access to penicillin (∼31 mg/kg) or VSL#3 probiotic (∼14 billion bacteria/mL) in drinking water. (**B**) Timeline of fecal material transplant (FMT) paradigm; B: baseline, amp: ampicillin, each dash = 1 day. (**C**) Hindpaw mechanical withdrawal thresholds of C57BL/6 mice during FMT paradigm. C57BL/6 mice received FMT from Townes AA (control) or Townes SS (SCD) mice; *N*=15-17. (**D**) Hindpaw withdrawal latency of C57BL/6 mice to dry ice application at FMT timepoint; unpaired t-test \*\*\*\**P*<0.0001. (**E**) Alpha diversity of fecal material collected from C57BL/6 mice prior to and following FMT. **(F**) Relative abundance of bacterial genera detected in fecal material collected from C57BL/6 mice prior to and following FMT. (**G**) Principal component analysis plot of weighted UniFrac beta diversity measure of fecal material at various timepoints in FMT paradigm. (**H**) Relative abundance of genus *Enterorhabdus* in control and SCD FMT recipients at FMT timepoint; LEfSe \**P*<0.05. (**I**) Relative abundance of genus *Roseburia* in control and SCD FMT recipients at FMT timepoint; LEfSe \**P*<0.05, n.d.: not detected. (**J**) Anatomical rendering of peripheral vagal innervation and neuronal cell body positioning in nodose jugular ganglia (NJG). (**K**) Representative firing patterns of NJG neurons isolated from control FMT or SCD FMT mice during step-wise depolarization (scale bar: 100 ms, 20 mV). (**L**) Spikes fired by NJG neurons during step-wise increasing pulses of depolarizing current; two-way RM ANOVA, main effect of current *P*<0.0001, current x genotype interaction *P*=0.08, 1X: rheobase, *n*=39-52 total cells isolated from 11 control or SCD FMT recipients at FMT timepoint). Bonferroni post hoc comparisons for all panels unless otherwise stated: \*\**P*<0.01, \*\*\**P*<0.001, \*\*\*\**P*<0.0001.

Fecal material transplant (FMT) experiments (Fig. 1B) were next performed to examine the extent to which the SCD microbiome induces pain in the absence of additional SCD pathology. After depleting the existing gut microbiome with antibiotics (Fig. S1), C57BL/6 mice that received SCD FMT developed mechanical allodynia that persisted for 3 weeks, whereas control FMT recipients demonstrated antibiotic-induced allodynia that was immediately reversed following FMT (Fig. 1C). SCD FMT recipients also developed cold allodynia (Fig. 1D) similar to what has been observed in individuals with SCD and SCD mouse models *(8,9)*, hypersensitivity to noxious punctate (Fig. S2A) and innocuous dynamic (Fig. S2B) mechanical stimulation, and, in male mice only, heat hypersensitivity (Fig. S2C). 16S rRNA gene sequencing was performed on feces collected from SCD and hemoglobin control FMT recipients prior to and 24 hr following the last FMT to determine if disease-related dysbiosis drives the divergent pain phenotypes. Overall, the FMT procedure allowed for recovery of antibiotic induced microbiome depletion to a similar extent in both SCD and hemoglobin control FMT recipients (Fig. 1E, 1F). Following FMT, feces from SCD and hemoglobin control FMT recipients contained similar bacterial populations (Fig. 1G, teal vs. gray circles). Only two bacterial genera – *Enterorhabdus* and *Roseburia* – were differentially abundant between treatment groups; both genera were present at higher levels in SCD FMT recipient feces (Fig. 1H, 1I). Notably, *Roseburia* is also elevated in feces collected from individuals with SCD *(2)*. Members of genus *Roseburia* are obligate anaerobes that produce butyrate, a short-chain fatty acid with well-documented anti-inflammatory effects *(10)* and the ability to induce fetal hemoglobin production *(11)*. Thus, it is unlikely that *Roseburia* is driving SCD FMT related pain, but rather, this microbe may be elevated in the SCD gastrointestinal tract as a compensatory mechanism to decrease disease-related inflammation and increase production of hemoglobin that is unable to polymerize.

Based on these analyses, it is likely that the genotype-specific pain behaviors observed following FMT do not result from the establishment of genetically diverse bacterial populations, at least not immediately following FMT. To determine if secondary effects of FMT on intestinal epithelium structural integrity explain the divergent pain phenotypes, gut leakiness was measured via FITC-dextran extravasation (Fig. S3A) and bacterial translocation (Fig. S3B). Both measures were similar between FMT genotypes. How then could the divergent FMT pain phenotypes be explained? An unexplored hypothesis is that FMT increased ascending sensory transmission from the gut. To test this idea, nodose/jugular ganglia (NJG), the structures in mice that contain vagal afferent cell bodies (Fig. 1J), were isolated from SCD and hemoglobin control FMT recipients 24 hr following the last FMT. NJG were then used for whole cell patch clamp recordings to measure intrinsic neuronal excitability. Two sub-types of NJG neurons became readily apparent during recordings: neurons that fired only once (1X; single-fire) upon sustained depolarization and neurons that fired repeatedly (>1X; multiple-fire) during prolonged current injection. Regardless of FMT genotype, single-fire neurons had more hyperpolarized resting membrane potentials, required significantly larger current injections to fire an action potential (*i*.*e*., higher rheobase), and had lower input resistance and action potential voltage thresholds relative to multiple-fire neurons (Table S1). The action potential half-width of single-fire neurons was also shorter than that of multiple-firers (Table S1). FMT genotype had no effect on any passive membrane property of single or multiple-fire neurons. However, when injected with depolarizing current steps, multiple-fire NJG neurons isolated from SCD FMT recipients fired more action potentials, on average, than neurons isolated from control FMT recipients (Fig. 1K, 1L). This increased excitability of multiple-fire NJG neurons may lead to enhanced glutamatergic signaling in interoceptive brain regions such as the nucleus of the solitary tract, which projects to neighboring brain structures important for pain sensation and perception (*e*.*g*., parabrachial nucleus) *(13)*.

### SCD associated heme catabolites in gastrointestinal tract induce persistent pain

Considering the lack of difference in bacterial populations between hemoglobin control and SCD FMT recipients, we hypothesized that FMT donor host or bacterial metabolites may be responsible for the SCD FMT-related increases in NJG neuronal excitability and pain behaviors. To this end, unbiased metabolomic screening was performed on SCD and hemoglobin control mouse donor feces as well as SCD and hemoglobin control FMT recipient feces. Of the >2,000 compounds identified across samples, heme catabolism products (Fig. 2A) were among the most dysregulated biochemicals between treatments.

**Fig. 2.**
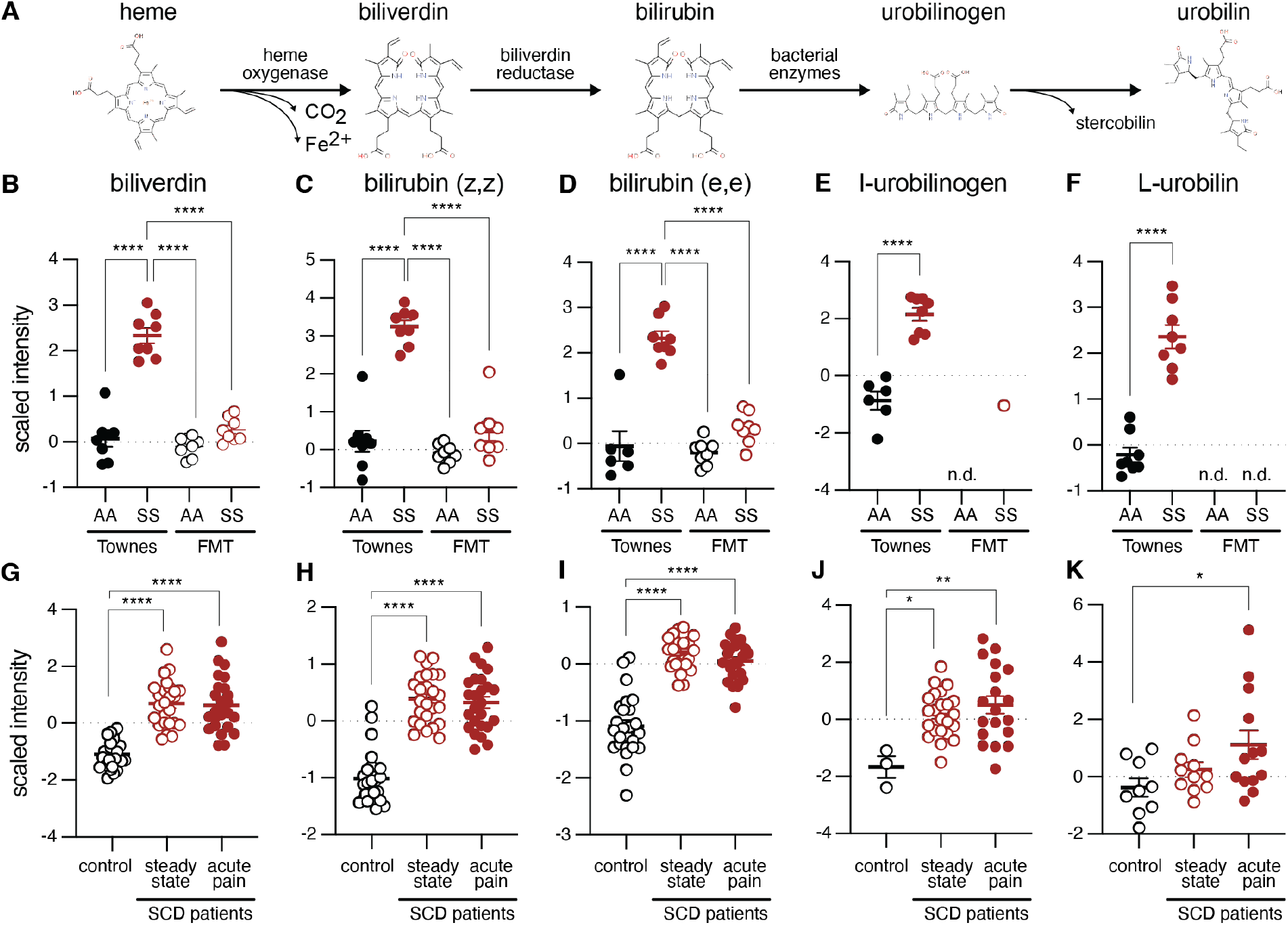
Heme breakdown products are elevated in patients and mouse models with SCD. (**A**) Heme catabolism pathway. Relative levels of (**B**) biliverdin, (**C**) bilirubin (z,z), **(D**) bilirubin (e,e), (**E**) I-urobilinogen, and (**F**) L-urobilin in fecal material from donor Townes AA, donor Townes SS, AA FMT recipient, and SS FMT recipient animals. Relative levels of (**G**) biliverdin, (**H**) bilirubin (z,z), **(I**) bilirubin (e,e), (**J**) I-urobilinogen, and (**K**) L-urobilin in plasma from patients with SCD at steady state and during an acute pain episode (same patients sampled at each timepoint) or from age, race-matched controls. Bonferroni post hoc comparison for all panels: \**P*<0.05, \*\**P*<0.01, \*\*\*\**P*<0.0001.

Biliverdin (Fig. 2B) and isomeric forms of bilirubin (Fig. 2C, 2D), two initial heme breakdown products, were elevated in fecal material from SCD mice relative to feces collected from hemoglobin control animals, control FMT recipients, and SCD FMT recipients. Urobilinogen, a compound generated when gut bacteria catabolize bilirubin, was also elevated in SCD fecal material relative to hemoglobin control feces and was only detected in one SCD FMT recipient sample (Fig. 2E); similar observations were also noted for urobilin (Fig. 2F).

We also measured heme catabolites from plasma of individuals with SCD since circulating metabolite signatures, particularly those generated by bacterial enzymes, are affected by gut dysbiosis (*14)*. Compared to race-matched controls, individuals with SCD had significantly elevated levels of biliverdin, bilirubin, and urobilinogen at steady state (*i*.*e*., at a regularly scheduled clinic visit with absence of health care utilization for pain for> 2 weeks prior to collection; Fig. 2G-I). There was no further increase in heme catabolite levels when the same patients were sampled during an acute pain episode (Fig. 2G-I). This contrasts with the traditionally held view that increased hemolysis occurs during vaso-occlusive events *(15)*. Thus, heme catabolism products may be the microbially-derived metabolites that drive chronic pain in individuals and mouse models with SCD. When recipient mice receive a SCD fecal transplant, they are receiving high concentrations of heme catabolites via oral administration. Given that recipients do not have ongoing hemolysis, they are able to breakdown the high levels of these compounds such that they excrete significantly less in their fecal material.

Since bilirubin is directly catabolized by gut bacteria, we hypothesized that it may be a critical component of the SCD FMT suspension that induces persistent pain behaviors and alters sensory neuron activity. To assess whether elevated bilirubin in the gastrointestinal tract induces widespread pain, hindpaw mechanical sensitivity was assessed following oral administration of bilirubin or vehicle control. Bilirubin-treated animals exhibited dose-dependent mechanical allodynia 30 min following administration suggesting an immediate nociceptive role for this compound (Fig. 3A).

**Fig. 3.**
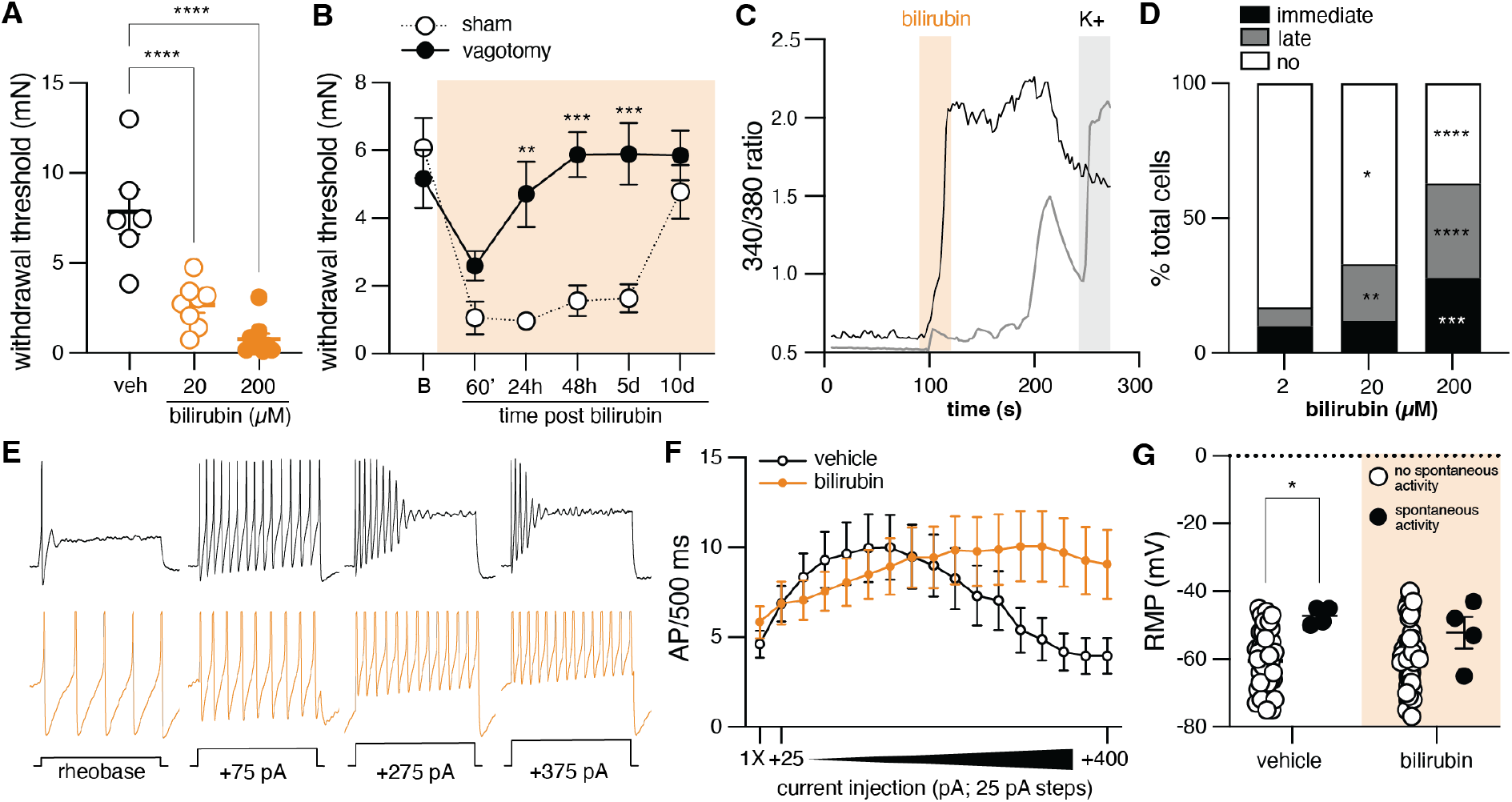
Oral bilirubin administration induces widespread, vagal nerve-dependent pain. (**A**) Hindpaw mechanical withdrawal thresholds of hemoglobin control mice 30 min following oral administration of bilirubin or vehicle. (**B**) Hindpaw mechanical withdrawal thresholds of hemoglobin control mice with intact (sham) or transected (vagotomy) vagus nerve following oral bilirubin (200 µM) administration; *N*=8-9. (**C**) Representative calcium traces from NJG neurons exposed to 30s pulse of bilirubin (200 µM) then KCl (50 mM). Immediate bilirubin responder shown in black; late bilirubin responder shown in gray. (**D**) Quantification of NJG neurons that exhibited immediate, late, or no response to 30s bilirubin exposure; *N*=129-185 neurons from 4-6 mice. (**E**) Representative firing patterns of NJG neurons isolated from vehicle or bilirubin (200 µM p.o.) treated mice during step-wise depolarization. (**F**) Spikes fired by NJG neurons during step-wise increasing pulses of depolarizing current; two-way RM ANOVA, main effect of current *P*<0.0001, current x genotype interaction *P*<0.0001, 1X: rheobase, *N*=18-23 total cells isolated from 7-8 mice 24 hr post treatment). (**G**) Resting membrane potential (RMP) of NJG neurons isolated from vehicle and bilirubin-treated mice; *N*=4 spontaneously active neurons per treatment, 41-46 not spontaneously active neurons per treatment. Bonferroni post hoc comparisons for all panels unless otherwise stated: \**P*<0.05, \*\**P*<0.01, \*\*\**P*<0.001, \*\*\*\**P*<0.0001.

Bilirubin-induced hypersensitivity persisted well beyond this timepoint; a single dose of bilirubin induced mechanical hypersensitivity that persisted for 5 days (Fig. 3B). To determine if this hypersensitivity resulted from bilirubin shifting the gut microbiome to more closely resemble that of SCD mice, 16S rRNA gene sequencing was performed on fecal material collected from SCD mice and hemoglobin control animals prior to, 24 hr following, and 5 days following oral bilirubin administration. Bacterial populations in SCD feces were more diverse than those detected in hemoglobin control samples collected before or after bilirubin treatment (Fig. S4A). Although bacterial populations changed in hemoglobin control animals following bilirubin administration, they remained distinct from those observed in SCD fecal material (Fig. S4B). Thus, SCD microbial populations result from additional selective pressures, not just elevated heme metabolites.

In the absence of significant bilirubin-induced shifts in the gut microbiota, we hypothesized that bilirubin induces widespread pain by altering ascending sensory neuron activity. Notably, bilirubin-induced mechanical hypersensitivity resolved 24 hr after administration in vagotomized mice (Fig. 3B) suggesting that sensory transmission through vagal afferents, the cell bodies of which reside in the NJG, is required for persistent bilirubin-induced pain. To assess the direct effects of bilirubin on sensory neuron function, calcium imaging was performed on NJG neurons isolated from naïve mice. Brief exposure to bilirubin induced immediate calcium flux in a subset of NJG neurons, induced a delayed (>30 s) response in a second subset of neurons, and failed to elicit any response in third subset of neurons (Fig. 3C). The proportion of bilirubin-responsive cells increased in a concentration-dependent manner (Fig. 3D).

To determine if bilirubin exposure also induces neuronal plasticity, whole cell patch clamp recordings were completed on NJG neurons isolated from animals that received oral administration of bilirubin or vehicle 24 prior to tissue collection. NJG neurons again clustered into two groups: single- and multiple-fire neurons (Table S2). No difference in any recorded measure was noted between single-fire neurons isolated from vehicle or bilirubin treated animals. Bilirubin did, however, alter the firing frequency of multiple-fire neurons (Fig. 3E, 3F); whereas NJG neurons from vehicle-treated animals exhibited use-dependent decreases in firing frequency, neurons isolated from bilirubin-treated mice fired with increasing frequency for the entirety of the stimulation protocol. Relative to vehicle control, bilirubin also hyperpolarized the resting membrane potential of spontaneously active neurons (Fig. 3G). In neurons isolated from vehicle-treated mice, the resting membrane potential of spontaneously active neurons was more depolarized than that recorded in neurons with no ongoing activity; no difference was noted in the resting membrane potential of spontaneously active and not-spontaneously active neurons isolated from bilirubin-treated mice. Based on these collective data, we propose that bilirubin is one of the primary metabolites that drives chronic SCD pain via activity on gut-innervating vagal afferents. Future experiments should determine if MRGPRA1 (MRGPRX4 in human) *(16)* or 5-HT_3_ *(17)*, two receptors expressed in rodent NJG *(18)* that are activated by bilirubin, underlie direct neuronal responses or plasticity resulting from bilirubin exposure. Results of those experiments may lead to the repurposing of approved drugs for use in SCD pain management (*e*.*g*., 5-HT_3_ antagonists).

### Chronic SCD pain is alleviated with *Akkermansia mucinophila* probiotic

Given that the SCD microbiome drives pain in the absence of additional SCD pathology, we next wanted to determine if chronic pain in SCD mice could be alleviated by manipulating their gut microbiome. To start, pain behaviors were measured in SCD mice after their gut microbiome was replaced with gut contents from healthy hemoglobin control mice or a separate cohort of SCD animals. SCD mice that received control FMT exhibited a transient, partial reversal of mechanical allodynia (Fig. 4A) and cold hypersensitivity (Fig. 4B) relative to SCD FMT recipients, but only for 24 hr following the final FMT.

**Fig. 4.**
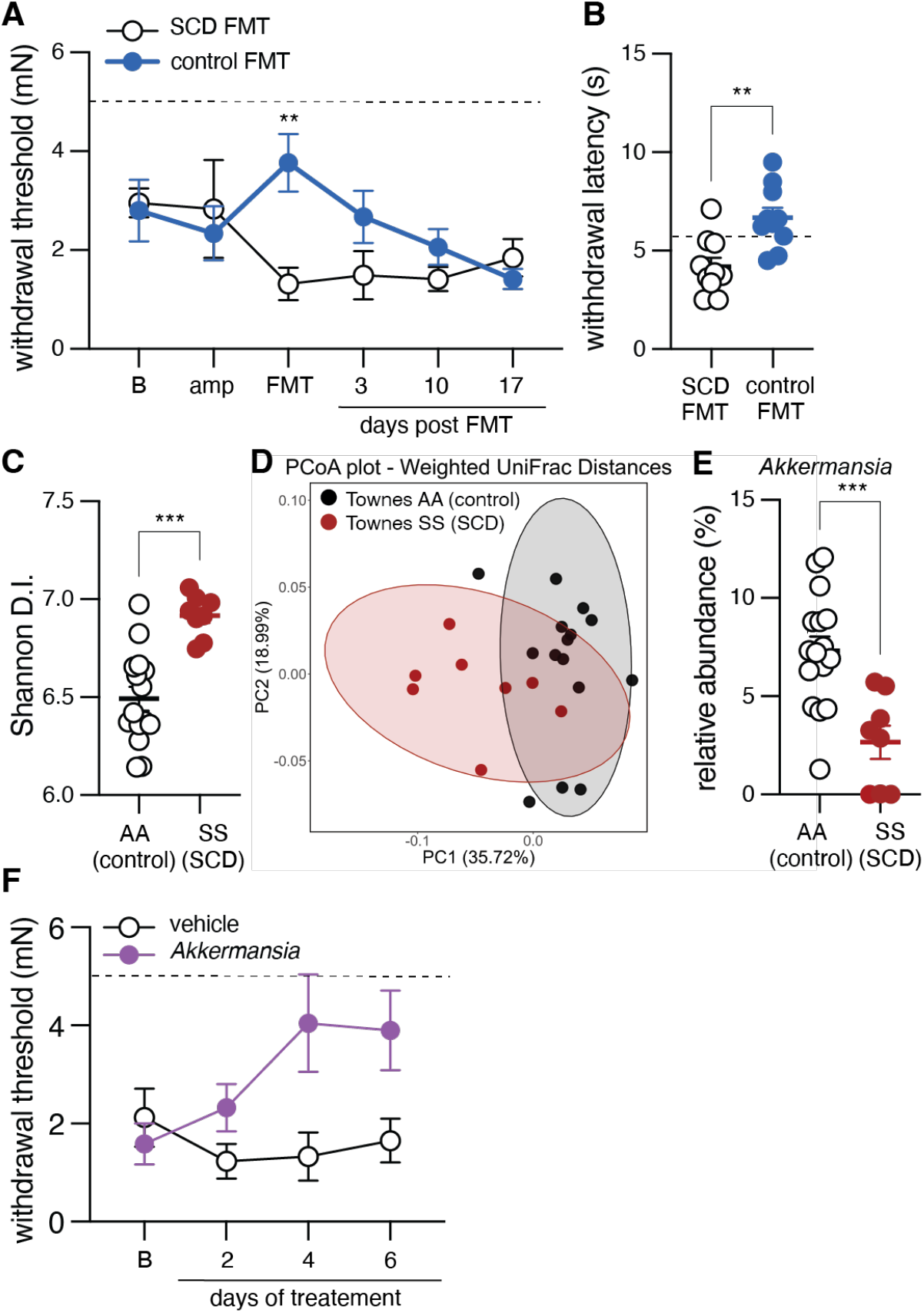
Identification of probiotics that alleviate chronic SCD pain. (**A**) Hindpaw mechanical withdrawal thresholds of Townes SS (SCD) mice during FMT paradigm. Mice received FMT from Townes AA (control) or separate cage of Townes SS (SCD) mice; dashed line indicates average withdrawal threshold of Townes AA control mice, B: baseline, amp: ampicillin timepoint, *N*=9-10. (**B**) Hindpaw withdrawal latency of Townes SS (SCD) mice to dry ice application at FMT timepoint; dashed line indicates average withdrawal latency of Townes AA control mice; unpaired t-test \*\**P*<0.01. (**C**) Alpha diversity of fecal material collected from Townes SS and AA mice. (**D**) Principal component analysis plot of weighted UniFrac beta diversity measure of fecal material from Townes SS and AA mice. (**E**) Relative abundance of genus *Akkermansia* in Townes SS and AA mice; unpaired t-test \*\*\**P*<0.001. (**F**) Hindpaw mechanical withdrawal thresholds of Townes SS mice prior to and following *Akkermansia* supplementation; two-way RM ANOVA main effect of treatment *P*<0.05, main time x treatment interaction *P*<0.05, dashed line indicates average withdrawal threshold of Townes AA control mice, B: baseline, *N*=6.

Based on these data, we concluded that control fecal material contained some bacteria or metabolite with analgesic properties, such that when introduced to the SCD gastrointestinal tract, persistent pain in these animals was alleviated. To this end, bacterial populations found in SCD and hemoglobin control fecal material were compared using 16S rRNA gene sequencing. Bacterial populations in SCD mouse feces had higher diversity (Fig. 4C) and were distinct from those found in hemoglobin control feces (Fig. 4D). Bacteria from several genera were present at different levels in SCD and control feces (Fig. S5). Most notably, bacteria from the genus *Akkermansia* were less abundant in feces from SCD mice (Fig. 4E), mirroring results obtained in feces collected from patients with SCD *(2). Akkermansia mucinophila* has recently gained attention as a promising next-generation probiotic due to its beneficial effects on gut epithelium integrity among other reasons *(19)*. To determine if *A. mucinophila* supplementation reversed chronic SCD pain, mechanical hypersensitivity was measured in SCD mice prior to and during daily *A. mucinophila* treatment. SCD mechanical allodynia was alleviated by *A. mucinophila* supplementation as soon as 4 days following the start of treatment (Fig. 4F).

In summary, these studies highlight the gut microbiome as a novel intervention site for chronic sickle cell disease pain management. Fecal content excreted by SCD mice induced persistent pain when orally administered to wildtype mice. In a similar fashion, bilirubin, a heme catabolite elevated in patients and mouse models of SCD, induced vagus nerve-dependent pain following oral administration. An association between bilirubin and SCD pain has only been suggested in one case report. After receiving a novel hemoglobin modifier, an adult man with SCD and jaundice reported no longer experiencing SCD pain and laboratory tests confirmed that he indeed had decreased circulating bilirubin levels (*20*). Given that both SCD FMT and oral bilirubin administration induced body wide pain, it is likely that these gastrointestinal signals altered pain signaling in brain regions that receive both interoceptive and nociceptive information. Deconstruction of these circuits will be the focus of future studies. It is notable that pathophysiological pressures in the gut resulting from SCD biology – namely hypoxia and byproducts of hemolysis – induce parallel shifts in the microbial populations that colonize both patients and transgenic mouse models with SCD. These similarities between species increase the translational potential of *Akkermansia mucinophila* as a novel analgesic for SCD pain management.

## Acknowledgments

The authors would like to thank the patients who donated clinical samples used in these studies and their family members, and members of the Stucky Lab – in particular, Tony Menzel – and the Center for Advanced Pain Studies at UT Dallas for their constant curiosity and constructive feedback on this project.

## Author contributions

Conceptualization: KES, AMB

Methodology: KES, SNA, VLE, TBW, MH

Investigation: KES, SNA, VLE, TBW, MH, DMR

Visualization: KES, SNA

Funding acquisition: KES, NHS, CLS, AMB

Supervision: KES, NHS, CLS, AMB

Writing – original draft: KES

Writing – review & editing: KES, SNA, VLE, TBW, MH, DMR, NHS, CLS, AMB

## Data and materials availability

All data are available in the main text or the supplementary materials.

## List of Supplementary Materials

Materials and Methods

Figs. S1 to S5

Tables S1 to S3

