## Supplemental Materials for "Gut microbiota and metabolites drive chronic sickle cell disease pain"

### Supplementary Materials

#### Materials and Methods

##### *Animals*

All animal protocols were in accordance with National Institute of Health guidelines and were approved by the Institutional Animal Care and Use Committees at the Medical College of Wisconsin (Milwaukee, WI; protocol #0383) and The University of Texas at Dallas (Richardson, TX; protocol #2022-0088). The following mouse strains were used throughout the paper:

*Townes transgenic mouse model of sickle cell disease (SCD)(4).* In this knock-in model, mouse α-globin genes are replaced with human α-globin genes, and the major and minor mouse β-globin genes are replaced with either: (1) normal human γ and human β-globin genes (hα/hα::$\beta^A/\beta^A$; "Townes AA", "hemoglobin control") or (2) normal human γ and sickle cell human β-globin genes (hα/hα::$\beta^S/\beta^S$; "Townes SS", "SCD"). Townes AA and SS mice were bred in house and maintained on a 14:10 light/dark cycle with *ad libitum* access to water and Teklad Global 16% Protein Rodent Diet chow (catalog #2916; irradiated). Equally sized cohorts of male and female mice were used in all experiments. Mice aged 8 -12 months were utilized in the FMT reversal experiments to capture the chronic phase of the disease phenotype; mice used in all other experiments were aged 3-9 months.

*C57BL/6 "wildtype" mice.* C57BL/6 mice were either bred in house or purchased from The Jackson Laboratory; purchased animals acclimated to housing facility for >7 days before experimental use. Purchased animals and animals bred in house were not intermixed, but rather used in independent experiments. C57BL/6 mice were maintained in the same housing facility and provided the same diet as Townes mice. Equally sized cohorts of male and female C57BL/6 mice aged 8-26 weeks were used.

##### *Animal treatments*

*Penicillin.* The antibiotic penicillin V was purchased from Sigma Aldrich and administered to animals in their drinking water for 7 days at a dose of ~31/mg/kg per day. This dose of penicillin is clinically relevant and was recently demonstrated to have long-term effects on the murine gut microbiome if administered early in life (*21)*. Penicillin-supplemented water was replaced every other day over the course of the experiment.

*Probiotics.* The probiotic VSL#3 is a mixture of eight strains of freeze-dried bacteria (*Streptococcus thermophilus, Bifidobacterium breve, Bifidobacterium longum, Bifidobacterium lactis, Lactobacillus acidophilus, Lactobacillus plantarum, Lactobacillus paracasei, Lactobacillus helveticus*). VSL#3 was purchased through Amazon (produced by Nutrilinea SRL, Varese, Italy; manufactured for Alfasigma USA, Inc., Covington, LA; lot 809158). VSL#3 was administered to animals in their drinking water for 7 days at a dose of ~14 billion bacteria/mL. This dose of VSL#3 was recently demonstrated to reverse visceral hypersensitivity in a rat model of colitis (*22*). VSL#3-supplemented water was replaced every day over the course of the experiment. *Akkermanisa* was purchased through Amazon (produced by Pendulum Therapeutics, San Francisco, USA; lot A22201001 and 02220117). *Akkermansia* as administered to animals in their drinking water for 7 days at a dose of ~ 20 million bacteria/mL.

*Fecal material transplant (FMT).* Ampicillin (0.5 g/L) was administered to animals in their drinking water for seven days. Fresh ampicillin-supplemented water was provided every other day, then replaced with normal drinking water on day 8. Twenty-four hours following ampicillin treatment cessation, animals received the first of three fecal transplants. Fecal transplants were prepared by dissolving freshly collected fecal material in 1X phosphate buffered saline, pH 7.4 (1 fecal pellet/1 mL of PBS); a mixture of fecal pellets was collected from 2-5 separate, unrelated cages of mice then vigorously vortexed to aid in pellet dissolution. Unfiltered fecal material suspension (200 µL/animal) was administered via oral gavage once daily for three days. Unless otherwise indicated, behavior was tested at the following FMT timepoints: "B" baseline, < 24 hr before initiating ampicillin treatment; "amp", day 3 or 4 following the start of ampicillin treatment; "FMT", the morning of the third FMT day (i.e., after 2 FMTs already performed).

*Bilirubin treatment.* Bilirubin was dissolved in vehicle containing 10% DMSO, 40% PEG 400, 5% Tween 80, 45% saline immediately prior to use. Bilirubin or vehicle solution (200 µL) was administered via oral gavage.

*Subdiaphragmatic vagotomy.* Nine days before behavior testing, C57BL/6 mice underwent bilateral subdiaphragmatic vagotomy or sham surgery. Briefly, once animals were anesthetized with isoflurane, a longitudinal incision was made through the skin and underlying fascia from the umbilicus region of the abdomen to just below the xiphoid process. A second incision of the same size was made in the underlying abdominal cavity wall. Using a sterile cotton-tipped swab, the internal organs were gently moved in order to visualize where the vagus nerve passes through the diaphragm. In vagotomized animals, the trunks of the left and right vagus nerve were cut immediately before ascending through the diaphragm; in sham animals, the nerve trunks were visualized, then the visceral organs were returned to their original positions. Both the abdominal cavity and skin incisions were secured with two sutures.

***Behavior tests***

Animals were habituated to testing chambers/apparatus for >1.5 hr prior to all behavior tests. The experimenter, who was blinded to treatment and/or genotype, remained in the behavior room for an additional 30 min prior to the start of behavior testing to allow for olfactory signal habituation. Animals were randomly assigned to treatment groups.

*von Frey punctate mechanical sensitivity testing.* Animals were placed into 10 x 10 x 15 $cm^3$ Plexiglass chambers on a raised 0.7 $cm^2$ wire platform. Calibrated monofilaments were delivered through the wire platform and applied to the plantar surface of each hindpaw following the up-down method (*23*); the 50% withdrawal threshold of each paw was calculated then averaged between paws (*24*). Toe flaring was not considered a "withdrawal".

*Plantar dry ice cold sensitivity testing.* Animals were placed into the aforementioned Plexiglass chambers on a raised 1/8" glass platform. Powdered dry ice was applied to the glass underneath the plantar surface of each hindpaw as previously described (*25*). The withdrawal latency of each paw was assessed five times, then averaged between paws.

*Noxious punctate mechanical sensitivity testing.* Animals were placed into the aforementioned Plexiglass chambers on a raised 0.7 $cm^2$ wire platform. A 25 Ga needle was applied to the center of the plantar surface of each hindpaw 10 times; response frequency and characterization were reported. Responses were characterized as follows: null (no paw withdrawal), normal (paw removed from wire mesh then immediately returned), or nocifensive (paw withdrawal accompanied by flicking, licking, biting, hovering of paw above mesh, slamming of paw back onto mesh surface).

*Dynamic light tough mechanical sensitivity testing.* Animals were placed into the aforementioned Plexiglass chambers on a raised 0.7 $cm^2$ wire platform. A fine, liner paintbrush (Princeton Good Synthetic Golden Taklon brush, size 2) was dragged – from heel to toes – across the glabrous surface of each hindpaw 10 times; response frequency and characterization was reported in a manner identical to needle testing.

*Radiant heat sensitivity testing.* Animals were placed into the aforementioned Plexiglass chambers on a raised 1/4" glass platform. A focal radiant heat source was applied to the glass immediately underneath the plantar surface of each hindpaw as previously described (*26*). The withdrawal latency of each paw was assessed five times, then averaged between paws.

***16s ribosomal RNA gene sequencing***

Animals were individually placed into glass beakers for fecal collection. For FMT-related sequencing, fecal material was collected from the same cohort of C57BL/6 mice at the following time points: < 24 hr prior to start of antibiotic administration (b: baseline), 4 days following the start of antibiotic administration (amp: ampicillin), and < 24 hr following the final of three FMTs. For bilirubin-related sequencing, fecal material was collected from Townes SS mice and the same cohort of Townes AA mice <24 hr before bilirubin administration, then again 24 hr and 5 days following oral administration of bilirubin (200 μM). Fecal material was collected immediately following excretion, placed into 0.5-1 mL of RNAlater, then frozen on dry ice. Fecal DNA was extracted using the Qiagen DNeasy Powerlyzer Powersoil Kit (cat. #12855-100) and the modified protocol described by Kommineni et al. (*27*). The V3/V4 region (341F – 806R) of the 16S rRNA gene was amplified via PCR and sequenced on the MiSeq platform (Illumina) using 2x300bp read technology at the University of Wisconsin-Madison Biotechnology Center, Madison, Wisconsin.

QIIME2 (v. 2022.8) was used to analyze the paired-end 16S rDNA sequencing reads (*28*). Sequences were imported and summarized to check quality. Cutadapt was used to trim primers from the reads (*29*).

Representative sequences were chosen using DADA2, which also removes chimeric (*30*). The representative sequences were then aligned (*31*), masked for hypervariable regions (*32*), and phylogenetic trees were produced (*33*). A classifier was generated to assign taxonomy to the reads using the 99% similarity files of the SILVA v. 138 and the 341-806 region (V3/V4) of the 16S gene (*34,35*). Taxonomy was assigned to the feature table to make taxonomy bar plots and to generate relative abundance tables.

Diversity metrics were run using the *core-metrics-phylogenetic* command of QIIME2. Alpha and beta diversity were analyzed using their respective commands, *alpha/beta-group-significance* (*36-38*). Alpha diversity metrics used a Kruskal-Wallis test to test for significance, while beta diversity metrics used a PERMANOVA test; both types of metrics used Benjamini-Hochberg multiple comparison tests. Principal Coordinate Analysis (PCoA) plots were examined using Emperor (*39, 40*) and finalized figures were made using the qiime2R package and ggplots2 in R (*41-43*). Differentially abundant taxonomy was calculated between two groups using LEfSe (Linear discriminate analysis (LDA) effect size) using default parameters except for -w being set to 1, as there were no subgroups to test (*44*). MaAsLin2 was used to determine differentially abundant taxa when multiple variables were being analyzed (*45*).

***Gut permeability***

Gut permeability was assessed using FITC-dextran. Twenty-four hours following the last of three FMTs, mice began an overnight fasting period. The following morning, Fluorescein isothiocyanate-dextran (FITC-dextran, Millipore Sigma, FD4; dissolved to concentration of 100 mg/mL in 1X phosphate buffered saline, pH 7.4) was orally administered to mice at a concentration of 44 mg/100 g body weight. Four hour later, trunk blood was collected via cardiac puncture. Blood was stored in the dark on ice, then centrifuged to separate plasma. Plasma was diluted 1:1 with 1X PBS, the mean fluorescence of each sample was detected on a 485 nm plate reader and circulating FITC-dextran levels were extrapolated from a standard curve.

***Bacterial translocation assessment*.**

Twenty-four hours following the last of three FMTs, mesenteric lymph nodes were collected from FMT recipient mice. Tissues were weighed and homogenized in 1X PBS (10 µL of PBS for every 1 mg of tissue). Serial dilutions of homogenate were grown on Trypticase soy agar, 5% sheep blood plates at 37°C; colony forming units were counted 48 hr later.

***Nodose/jugular ganglia (NJG) calcium imaging and electrophysiology***

*Nodose/jugular ganglia neuronal isolation and culture.* Approximately 24 hours following the final FMT or following oral bilirubin (200 µM) administration, bilateral nodose/jugular ganglia (NJG; contained within the same ganglion in mouse) were isolated from C57BL/6 wildtype mice. After isoflurane anesthesia and cervical dislocation, an incision was made in the lateral aspect of the neck at the level of the hyoid bone. Following removal of the sternocleidomastoid and masseter muscles, the facial (VII), glossopharyngeal (IX), vagus (X), accessory (XI), and hypoglossal (XII) cranial nerves were visible. After identifying the vagus nerve by observing its parallel orientation to the carotid artery, the nerve trunk was traced in a rostral fashion until it entered the skull at the jugular foramen; the NJG appeared as white swellings within each jugular foramen. Both NJG were excised, incubated in DMEM/Ham's F12 medium containing 10 mg/mL collagenase for 40 min at 37°C, then 0.5% trypsin for 45 min at 37°C. Following trypsin neutralization in 3% heat-inactivated horse serum, NJG were mechanically dissociated then plated on laminin-coated glass coverslips. Neurons were incubated overnight at 37°C, 5% $CO_2$ in DMEM/Ham's F12 medium supplemented with 10% heat-inactivated horse serum, 2 mM L-glutamine, 1% glucose, 100 units/mL penicillin, and 100 µg/mL streptomycin. Patch clamp electrophysiology was performed on neurons 12-24 hr following tissue isolation.

*Whole-cell patch clamp electrophysiology.* Patch clamp recordings of isolated NJG neurons were performed following overnight culture. Neurons were visualized using a Nikon Eclipse TE200 inverted microscope, and coverslips were continuously superfused with extracellular buffer (140 mM NaCl, 2.8 mM KCl, 2 mM $CaCl_2$, 1 mM $MgCl_2$, 10 mM HEPES, 10 mM glucose, and 8.8 mM sucrose, pH 7.4 ± 0.02 and 310 ± 3 mOsm). Borosillicate glass pipettes (2.4 to 4.2 MΩ) filled with internal solution (135 KCl, 4.1 $MgCl_2$, 2 EGTA, 0.2 mM NaGTP, 2.5 mM $ATPNa_2$, and 10 mM HEPES, pH 7.2 ± 0.02 and 290 ± 2 mOsm) were pulled using a Sutter Instruments P87 pipette puller and used to perform patch clamp recordings. Series resistance was maintained at <10 MΩ and compensated at 60%. Neuronal capacitance was fully compensated and continuously monitored to ensure stable recording conditions.

Whole-cell recordings were obtained in current-clamp mode using a HEKA EPC10 amplifier, and recordings were obtained using Patchmaster software. Data were collected from neurons with a resting membrane potential (RMP) ≤ -40mV. In FMT recordings, neurons were held at -70mV during; in bilirubin recordings, neurons were held at RMP. Measures of intrinsic excitability were recorded using the following protocols: (1) Voltage-current (*V-I*) relations were obtained using 20 sweeps of 500 ms ascending current pulses (5 pA stepwise increase from holding current). The plateau voltage deflection was plotted against current amplitude, and input resistance was determined from the slope of the *V-I* plot where voltage sweeps did not exhibit active conductance. (2) Action potential (AP) properties were measured using an ascending series of 5 ms depolarizing current pulses. Rheobase was defined as the first current to elicit a single spike. AP threshold was determined from a derivative function, where dV/dt first exceeded 28 mV/ms. AP amplitude was determined relative to AP threshold, and AP half-width was measured as the width at half of the AP amplitude. (3) A series of depolarizing current steps were used to elicit action potentials. In FMT recordings, nine 500 ms (range: rheobase to 1600 pA above rheobase; 200 pA increments, 20 s intervals) were used; in bilirubin recordings, seventeen 500 ms (range: rheobase to 400 pA above rheobase; 25 pA increments, 20 s intervals) were used. (4) The postburst afterhyperpolarization (AHP; avg. of 3 sweeps at 20 s intervals) was examined using a 50 Hz burst of 10 spikes evoked by a 2 ms suprathreshold current pulse. The average AHP amplitude during the first 150 ms following current offset was used to determine the medium AHP (mAHP), while the AHP amplitude at 1 s following current offset was used to determine the slow AHP (sAHP). Recorded neurons were classified as either single-fire (firing only 1 AP regardless of current magnitude) or multiple-fire (firing more than 1 AP at suprathreshold currents). Spontaneous activity was recorded in current clamp mode for 2 min after voltage-current relationships were assessed and prior to depolarizing current injection steps.

*NJG neuronal calcium imaging.* Calcium imaging of isolated NJG neurons from naïve, C57BL/6 mice was performed following overnight culture. Neurons were loaded with the dual-wavelength, ratiomtic calcium indicator dye, Fura-2-AM (2.5 µg/mL in 2% bovine serum albumin) for 45 min, then washed with extracellular buffer (150 mM NaCl, 5.6 mM KCl, 2 mM $CaCl_2$, 1 mM $MgCl_2$, 10 mM HEPES, 8 mM glucose, pH 7.4 ± 0.02 and 320 ± 3 mOsm) for 30 min. Coverslips were mounted onto a Nikon Eclipse TE200 inverted microscope, and superfused with extracellular buffer at a rate of 6 mL/min. Cells were superfused with bilirubin (2, 20, or 200 µM) prepared in 10% DMSO, 40% PEG 400, 5% Tween 80, 45% PBS for 30 s, then 50 mM KCl for 30s. Bilirubin was prepared immediately before use and protected from light exposure. Fluorescence images were captured at 340 and 380 nm using NIS Elements Software. Immediate responders were cells that exhibited a ≥ 20% increase in 340/380 nm ratio from baseline during the 30 s in which bilirubin was superfusing cells; late responders were cells that exhibited a ≥ 20% increase in 340/380 nm ratio from baseline after bilirubin superfusion ended, but before 50 mM KCl exposure occurred. Null responders were cells that only exhibited a ≥ 20% increase in 340/380 nm ratio from baseline during 50 mM KCl exposure.

***Metabolite screens***

*Mouse fecal material.* Unbiased fecal metabolite screening was performed by Metabolon (Morrisville, NC). Briefly, fecal material was collected from FMT donors and recipients using the methods described for 16s sequencing. Upon excretion, fecal material was immediately collected and transferred to an empty, pre-chilled tube. Samples were frozen on dry ice, and shipped to Metabolon where they were prepared using the automated MicroLab STAR system (Hamilton Company). Sample compounds were measured using a Waters ACQUITY ultra-performance liquid chromatography and Thermo Scientific Q-Exactive high resolution/accurate mass spectrometer interfaced with a heated electrospray ionization (HESI-II) source and Orbitrap mass analyzer operated at 35,000 mass resolution. Compounds were identified by comparison to purified standards, and relative metabolite concentrations were calculated using Metabolon hardware and software.

*Patient plasma metabolomic screen*. The study was approved by the Institutional Review Board (IRB) at Children's Wisconsin. Informed written consent was obtained from the participants' legal guardian and assent was obtained from the child when age appropriate. Study participants included male and female children aged 7-19 years with SCD and healthy Black controls. Patient demographics are described in Table S3. Patients were recruited during routine visits to the sickle cell clinic or when hospitalized for acute pain. Plasma was collected during baseline state of health, defined as the absence of an acute care visit for pain or another sickle cell disease complication for at least two weeks, or during hospitalization for acute pain. Plasma was collected by peripheral venipuncture, immediately centrifuged and stored in 500 µl aliquots in -80°C freezer until use.

Deidentified patient samples were sent to Metabolon for the same unbiased metabolomic screening performed on mouse fecal material.

***Statistical analysis***

Individual data points were presented whenever possible. All other data are presented as group means ± SEM. Unless otherwise described, data were analyzed using GraphPad Prism 9; results were considered statistically significant when $P < 0.05$. Data from each sex was independently analyzed then combined if no sex difference was observed. Sex differences were observed in heat sensitivity following SCD FMT; data for each sex are presented in Fig S2C, S2D.

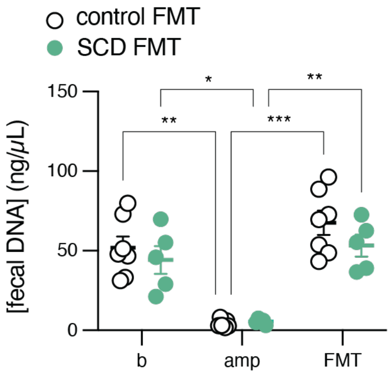


**Fig. S1**. **DNA concentration in fecal material collected from mice throughout FMT paradigm**. 2-way RM ANOVA main effect of time $P<0.0001$; Bonferroni post-hoc tests: *$P<0.05$, **$P<0.01$, ***$P<0.001$; b: baseline, amp: ampicillin, FMT: fecal material transplant).

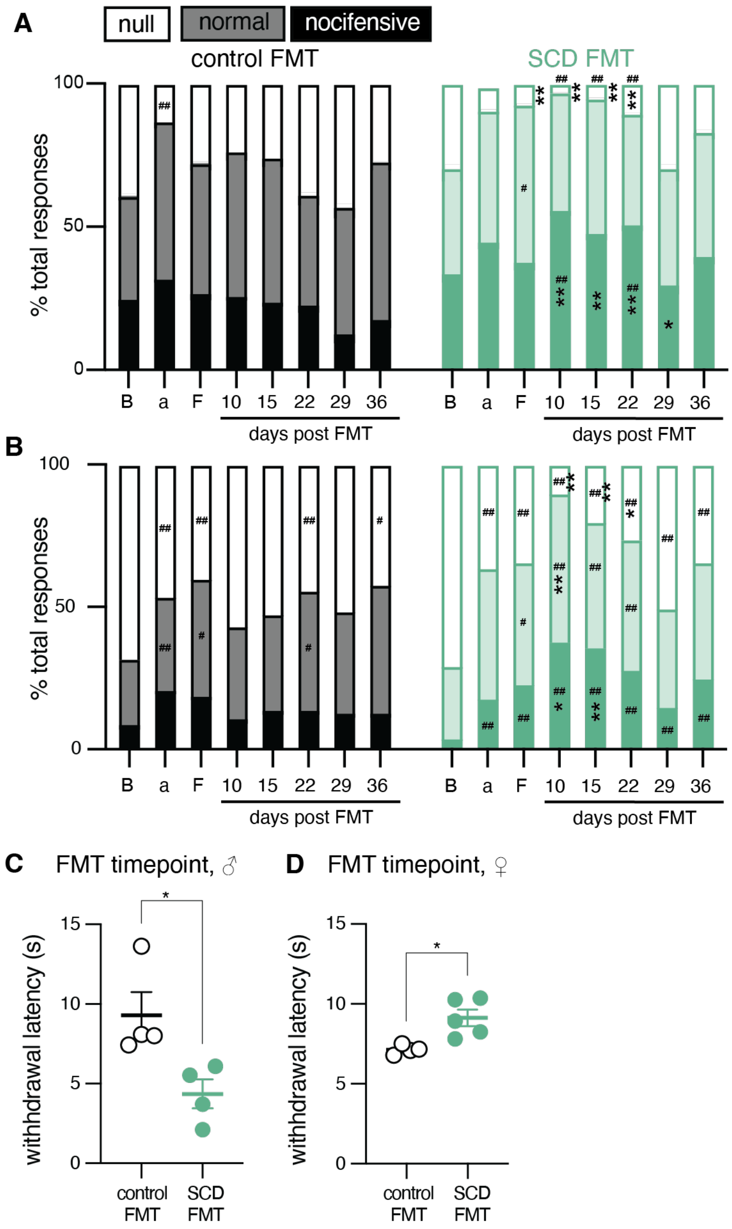


**Fig. S2. SCD FMT induces persistent pain in recipients**. (**A**) Noxious needle responses of hemoglobin control and SCD FMT recipients throughout FMT paradigm; Fisher's exact post hoc comparisons: *$P$<0.05 control vs. SCD, **$P$<0.01, control vs. SCD, #$P$<0.05 timepoint vs. baseline, ##$P$<0.01, timepoint vs. baseline, $N$=8-11. (**B**) Dynamic paintbrush responses of hemoglobin control and SCD FMT recipients throughout FMT paradigm; Fisher's exact post hoc comparisons: *$P$<0.05 control vs. SCD, **$P$<0.01, control vs. SCD, #$P$<0.05 timepoint vs. baseline, ##$P$<0.01, timepoint vs. baseline, $N$=8-11. (**C**) Hindpaw withdrawal latency of male mice to radiant heat application at FMT timepoint; Mann-Whitney test *$P$=0.0286. (**D**) Hindpaw withdrawal latency of female mice to radiant heat application at FMT timepoint; Mann-Whitney test *$P$=0.0159.

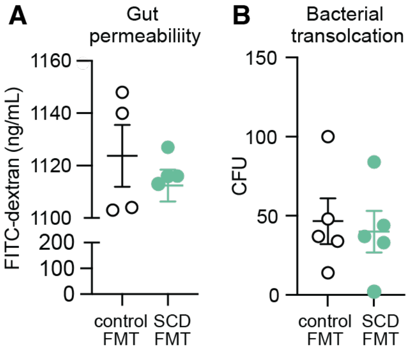


**Fig. S3. SCD FMT does not induce gross changes in gastrointestinal tract integrity.** (**A**) Serum levels of FITC-dextran in control and SCD FMT recipients at FMT timepoint. (**B)** Colony forming units present on TSA III Blood Agar after plating homogenized mesenteric lymph node tissues from control and SCD FMT recipients at FMT timepoint.

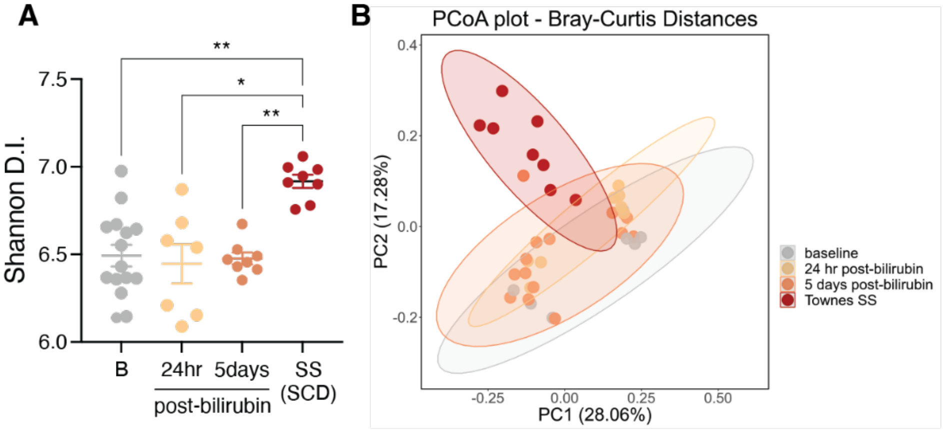


**Fig. S4. Oral bilirubin treatment alters gut microbiota. (A)** Alpha diversity of fecal material collected from hemoglobin control mice before and at various time points following oral bilirubin (200 μM) administration as compared to SCD mice fecal material (Shannon diversity index, $P$=0.00123). (**B)** Beta diversity observed in fecal material collected from hemoglobin control mice before and at various time points following oral bilirubin administration (200 μM) as compared to SCD mice fecal material (Bray-Curtis dissimilarity, $P$=0.001).

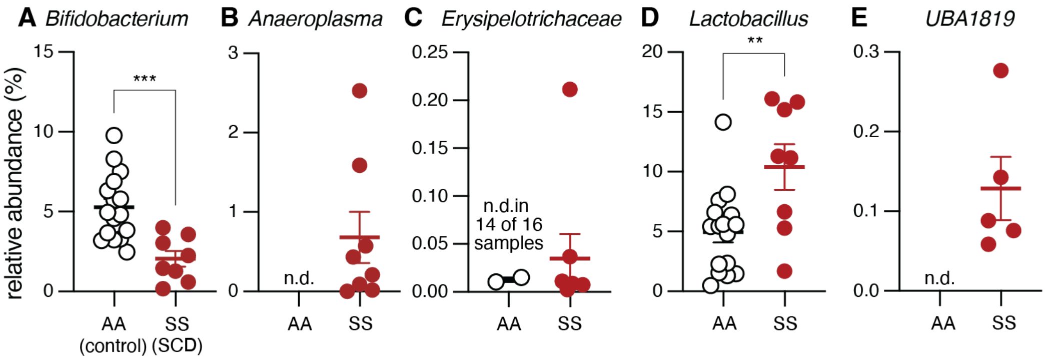


**Fig. S5: Differentially abundant bacteria in SCD and hemoglobin control fecal material.** MaAsLin2 multivariate analysis of 16S rDNA sequencing of Townes SS (SCD) and Townes AA (control) fecal material revealed significant differences in the relative abundance of the following bacteria: (**A**) *Bifidobacterium*, (**B**) *Anaeroplasma*, (**C**) *Erysipelotrichaceae*, (**D**) *Lactobacillus*, (**E**) *UBA1819*; Kruskal-Wallis test ***P*<0.01, ****P*<0.001.

| | control FMT | | SCD FMT | |
|---|---|---|---|---|
| | ***1X*** | ***>1X*** | ***1X*** | ***>1X*** |
| cell number | 30 | 39 | 33 | 52 |
| cell diameter (µm) | 27.09 ± 0.62 | 25.72 ± 0.59 | 27.91 ± 0.65 | 27.08 ± 0.43 |
| capacitance (pF) | 37.50 ± 3.28 | 32.37 ± 2.23 | 34.28 ± 1.97 | 34.17 ± 2.23 |
| RMP (mV) [a] | -66.16 ± 1.78 | -62.56 ± 2.18 | -67.00 ± 1.48 | -60.74 ± 1.74 |
| rheobase (pA) [a] | 797.9 ± 99.46 | 240.5 ± 18.18 | 923.6 ± 93.82 | 283.4 ± 26.98 |
| input resistance (mΩ) [a] | 153.9 ± 25.04 | 369.8 ± 35.44 | 144.4 ± 26.35 | 372.2 ± 27.90 |
| AP threshold (mV) [a] | -29.91 ± 1.25 | -21.09 ± 1.26 | -27.38 ± 1.61 | -22.15 ± 0.89 |
| AP amplitude (mV) [b] | 64.12 ± 2.18 | 61.34 ± 1.47 | 61.05 ± 2.00 | 65.10 ± 1.11 |
| AP half-width (ms) [a] | 1.38 ± 0.09 | 1.73 ± 0.06 | 1.21 ± 0.07 | 1.80 ± 0.06 |

**Table S1. Passive and active membrane properties of NJG neurons isolated from C57BL/6 mice 24 hours following SCD or control FMT**. Mean ± SEM listed for all properties. 2-way ANOVA main effect of fire type: [a]; 2-way ANOVA interaction between fire type x treatment: [b]; Bonferroni post-hoc tests: **P*<0.05, ***P*<0.01; 1x: fires only once, >1X: fires more than once, RMP: resting membrane potential, AP: action potential).

| | vehicle | | bilirubin | |
|---|---|---|---|---|
| | *1X* | *>1X* | *1X* | *>1X* |
| cell number | 22 | 23 | 31 | 18 |
| cell diameter (µm) [a,b] | 21.63 ± 1.02* | 18.71 ± 0.48* | 19.03 ± 0.65 | 18.29 ± 0.72 |
| capacitance (pF) [a] | 50.22 ± 10.88 | 22.96 ± 2.08 | 35.62 ± 7.35 | 22.46 ± 2.68 |
| RMP (mV) [a] | -62.14 ± 1.81 | -56.78 ± 1.89 | -59.61 ± 1.81 | -56.94 ± 2.14 |
| input resistance (mΩ) [a] | 367.6 ± 81.67 | 756.5 ± 165.3 | 388.1 ± 56.18 | 860.5 ± 119.9 |
| rheobase (pA) [a] | 523.6 ± 130.0 | 82.61 ± 14.26 | 701.6 ± 212.0* | 76.11 ± 12.34* |
| AP threshold (mV) [a] | -25.21 ± 3.46 | -16.11 ± 2.55 | -25.38 ± 2.04* | -13.46 ± 2.78* |
| AP amplitude (mV) [a] | 75.96 ± 5.72 | 90.33 ± 3.37 | 67.76 ± 4.88** | 92.22 ± 4.03** |
| AP half-width (ms) | 2.97 ± 0.51 | 2.72 ± 0.28 | 3.28 ± 0.38 | 2.75 ± 0.30 |
| spontaneous activity | 4 of 23 | | 4 of 18 | |

**Table S2. Passive and active membrane properties of NJG neurons isolated from bilirubin (200 µM) or vehicle treated mice**. Mean ± SEM listed for all properties. 2-way ANOVA main effect of fire type: [a]; 2-way ANOVA main effect of treatment: [b]; Bonferroni post-hoc tests: *$P$<0.05, **$P$<0.01; 1x: fires only once, >1X: fires more than once, RMP: resting membrane potential, AP: action potential.

| | Controls | SCD patients |
|---|---|---|
| Number of participants | 25 | 25 |
| Age (years; mean ± SD) | 10.76 ± 3.49 | 12.84 ± 2.66 |
| Gender: female | 14 (56%) | 13 (52%) |
| *Genotype* | | |
| HbSS | N/A | 17 (68%) |
| HbSC | | 6 (24%) |
| HbSβ+thal | | 1 (4%) |
| SO Arab | | 1 (4%) |

**Table S3. Subject demographics and clinical characteristics.**
